# FUSE-PhyloTree: Linking functions and sequence conservation modules of a protein family through phylogenomic analysis

**DOI:** 10.1101/2025.03.28.645931

**Authors:** Olivier Dennler, Elisa Chenel, François Coste, Samuel Blanquart, Catherine Belleannée, Nathalie Théret

## Abstract

FUSE-PhyloTree is a phylogenomic analysis software for identifying local sequence conservation associated with the different functions of a multi-functional (e.g., paralogous or multi-domain) protein family. FUSE-PhyloTree introduces an original approach that combines advanced sequence analysis with phylogenetic methods. First, local sequence conservation modules within the family are identified using partial local multiple sequence alignment. Next, the evolution of the detected modules and known protein functions is inferred within the family’s phylogenetic tree using three-level phylogenetic reconciliation and ancestral state reconstruction. As a result, FUSE-PhyloTree provides a gene tree annotated with both predicted sequence modules and ancestral gene functions, enabling the association of functions with specific sequence regions based on their co-emergence.

**Availability and Implementation:** FUSE-PhyloTree is provided as Docker and Singularity images including all the required software tools. Images, source code, test data, and documentation are available at https://github.com/OcMalde/fuse-phylotree.

**Supplementary Information:** An illustration of the application of FUSE-PhyloTree to the fibulin protein family is presented in the Appendix.

## Introduction

FUSE-PhyloTree is dedicated to the study of multifunctional protein families, including for instance paralogous or multi-domain proteins. These families, containing proteins with different and possibly multiple functions, are challenging to study experimentally and are often incompletely annotated, with missing functional annotations for some proteins and limited identification of the functionally important sequence regions. While common, these families are difficult to analyze with traditional computational tools due to their complex evolution, which can involve sequence duplication, insertion, and deletion events, along with varying selection pressures on specific sequence regions as functions are gained or lost.

Methods for refining protein families into functional families (FunFams) and predict their functional sites are reviewed by Rauer et al. (11). The first approach relies on detecting known domains (12), such as those from the integrated database Interpro (10). However, the number of known functional domains is limited, and the precision of the predicted annotations can be insufficient for the family functional characterization. Rauer et al. (11) highlights thus two main approaches: (i) constructing sequence similarity network, typically by pairwise comparisons, and (ii) clustering the multiple sequence alignment (MSA) of the family with respect to a phylogenetic tree.

The advantage of the alignment approach (ii) over approach (i) is to consider the entire family sequence information. Yet, proteins within multi-functional families have distinct functions supported by different sequence regions. They are thus difficult to characterize with classical MSA since functionally important regions may be conserved only within specific subsets of the family and may occur at different locations. To deal with such families, the first originality of FUSE-PhyloTree lies in employing an original alignment method developed by our team (2, 7): the *partial local multiple alignment* (PLMA). A PLMA identifies all compatible blocks of local sequence conservation shared by at least two sequences, each block consisting in a local alignment of significantly similar segments from some of the sequences to align. These blocks, referred to as sequence conservation modules or simply *modules*, form the basis of our approach.

According to the definition of a PLMA block, identifying a set of modules that encompass all key functional sequence segments requires that each function of interest is represented by at least two sequences in the aligned set. This can be achieved by incorporating orthologs within the family. However, the association of the specific (sets of) modules with the particular functions will still remain unknown. The second originality of FUSE-PhyloTree lies in combining this sequence analysis with a phylogenomic approach (4). While approach (ii) uses phylogenetic inference to cluster sequence alignment, FUSE-PhyloTree proposes to use advanced methods to infer and map the *evolutionary history* of both the *functions* and the *modules* onto the phylogenetic tree of the protein family (which can include paralogs and orthologs). Concerning the functions, FUSE-PhyloTree uses the maximum likelihood method for ancestral state reconstruction proposed by Ishikawa et al. (6) to infer their presence/absence at each ancestral node of the gene tree. With respect to the sequences, estimating the modules’ history is much more complex, as gene regions undergo their own events (gain/loss) independently of their host gene’s history, as well as the genes encounter their own events (paralogous duplication/loss) independently of their host species’ history. To map the modules’ history onto the gene tree, FUSE-PhyloTree employs the reconciliation method by Li and Bansal (9), which accounts for duplication, transfer, and loss events occurring during both the modules’ evolution and the genes’ evolution.

As a result, FUSE-PhyloTree generates a phylogenetic tree annotated with both the ancestral function and module content of its nodes, enabling the study of their associations. Natural selection preserves the protein sequence regions that are critical for essential functions, and the conservation of functions and modules since a given common ancestor suggests their potential association. By identifying the co-emergence of functions and modules within the annotated gene tree, FUSE-PhyloTree proposes candidate module sets likely to be linked to specific functions. Additionally, FUSE-PhyloTree enables interactive visual exploration of the annotated gene tree with iTOL (8), enabling in-depth examination of the co-evolution of functions and modules.

FUSE-PhyloTree prototype was initially developed to support research on the ADAMTS/TSL family (2). In the supplementary data, we present an illustration of the FUSE-PhyloTree application to another protein family, the fibulins (Supplementary Appendix), along with the files required to reproduce the experiment (See GitHub for Files).

## FUSE-PhyloTree workflow

Given a multi-functional protein family, FUSE-PhyloTree integrates sequence and phylogenomic analyses to predict functional regions by associating conserved sequence regions with specific functional annotations. We present here the different steps of FUSE-PhyloTree’s pipeline (Fig. 1).

**Fig. 1.**
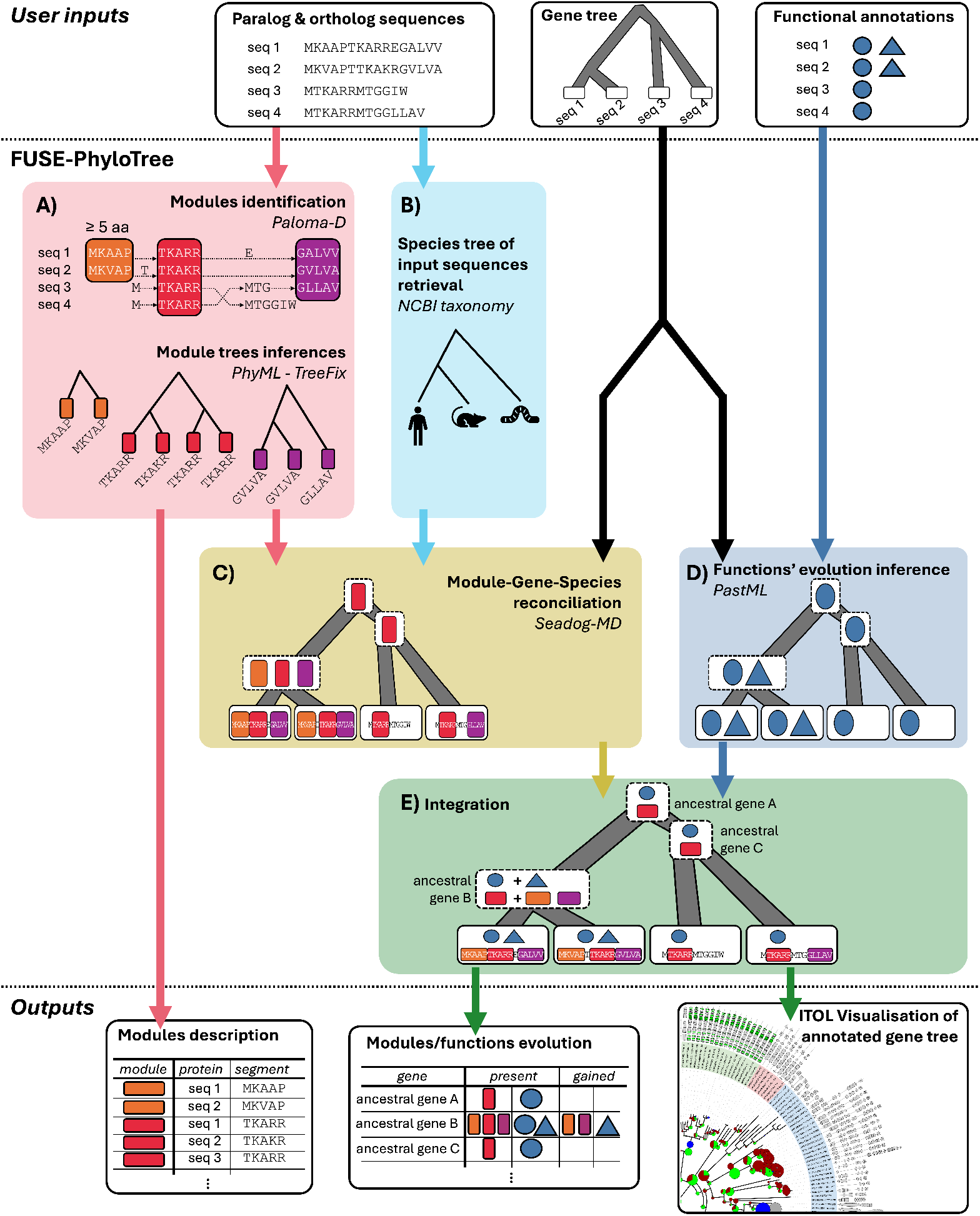
FUSE-PhyloTree workflow. FUSE-PhyloTree takes three input data files: a set of protein sequences (FASTA file), a binary rooted gene tree for these sequences (newick file), and their associated functional annotations (e.g. protein-protein interactions, CSV file). In this schema, there is only two functions symbolized by a blue triangle and a blue circle respectively. A) Identification of sequence conservation modules using Paloma-D. For each module, a phylogenetic module tree is computed using PhyML and TreeFix considering the gene tree as a reference. The orange, red and magenta colored boxes indicate the modules, defined by aligned protein segments (top of the panel) whose phylogenies are inferred (bottom). B) The species tree corresponding to the input sequences is retrieved from NCBI taxonomy. C) The species, gene and module trees are reconciled using SEADOG-MD, to infer the presence of modules along the gene tree. D) The evolution of functions along the gene tree is inferred using PastML, considering the gene tree and the protein functional annotations as input and mapping the presence of functions at each tree node. E) The integration of previous steps yields a gene tree annotated with the presence of both modules and functions, facilitating the exploration of sequence-function associations through their evolutionary history. Output files include interactive iTOL representations (text files - iTOL) and tabular files listing the presence, gain, and loss of modules and functions at each gene node.

FUSE-PhyloTree takes as input the three following files: 1) a FASTA file containing protein sequences of the target family (including both paralogs and orthologs) with the indication of their NCBI taxonomic identifier (*taxid*), 2) a newick file for the rooted binary gene tree corresponding to these sequences, and 3) a tabular file (CSV format) of functional annotations of interest of proteins, for instance their identified protein-protein interactions.

FUSE-PhyloTree method involves five key steps: A) Sequence conservation modules are identified using Paloma-D (2, 7), and each module’s independent evolutionary history is inferred as a phylogenetic module tree with PhyML (5), corrected using TreeFix (13) considering the gene tree as template. B) The species tree for the input sequences is retrieved from NCBI taxonomy using their *taxid*. C) Module, gene, and species trees are reconciled using SEADOG-MD (9) to infer modules evolution, identifying which modules are present at each ancestral gene node. D) Simultaneously, the evolution of functions in the gene tree is inferred using PastML (6), mapping which functions are present at each ancestral gene node. E) Finally, modules and functions’ evolution are integrated on the gene tree, producing a unified scenario that details the evolution of both sequence conservation modules and functions—mapping their presence, gain, and loss at each ancestral gene node.

FUSE-PhyloTree outputs consist in the detailed descriptions of the sequence conservation modules and the annotated gene tree, showing the presence, gain, and loss of these modules and functions. To reveal potential sequence-function relationships, FUSE-PhyloTree proposes associations between sequence regions and functions based on their simultaneous emergence during gene evolution. Specifically, the outputs consist of: i) a tabular file listing each module, the sequences where they are found, their positions and corresponding segments, ii) a tabular file mapping the presence, gain, and loss of modules and functions at different gene nodes on the gene tree, and iii) all files for iTOL (8) interactive visualizations of all generated data. Additional data from intermediate steps, such as the PLMA computed by Paloma-D and all probabilities of function presence at various gene nodes computed by PastML, are available in a directory for further analysis if required. A full list of the generated files and their contents is detailed in the GitHub documentation.

Additionally, FUSE-PhyloTree provides two optional utility steps that users may perform prior to the main analysis: i) Homologous sequence retrieval: users can expand their initial protein set by retrieving homologous sequences from a predefined set of nine reference species—including human and other animals—using RefSeq accession numbers and precomputed orthogroups. For each gene, its longest isoform is then selected, preparing a formatted sequence input ready for the FUSE-PhyloTree workflow. ii) Gene tree inference: as an alternative to providing a gene tree, FUSE-PhyloTree can automatically infer one directly from the input sequences, utilizing standard phylogenetic tools (MUSCLE, PhyML). However, caution is advised regarding the inferred tree’s root, as the accuracy of the gene tree topology is critical for FUSE-PhyloTree analysis. For more advanced usage, users can also provide their own species tree, PLMA, or even all intermediate files prior to the final integration step. See the GitHub documentation for instructions on usage and the appendix for an illustration of the application of FUSE-PhyloTree on a real example.

## Implementation

FUSE-PhyloTree is implemented as a modular pipeline that integrates multiple standalone software tools with custom Python scripts, handling tasks from input/output formatting to newly developed algorithms for integrating their outputs.

FUSE-PhyloTree relies on the following standalone software: MUSCLE (3) for multiple sequence alignment (MSA), trimAl (1) for MSA trimming, PhyML (5) for phylogenetic tree inference, TreeFix (13) for tree correction, Paloma-D (2, 7) for PLMA computation, SEADOG-MD (9) for three-level DGS (Domain-Gene-Species) reconciliation, and PastML (6) for ancestral scenario reconstruction of functions.

For standard use, FUSE-PhyloTree is provided as Docker and Singularity images, bundling all required software, dependencies, and scripts for easy use across Linux, Windows and MacOS systems and high performance computing clusters. This setup allows users to run the pipeline with minimal configuration, while all custom Python scripts are freely available in the GitHub repository.

## Availability

The FUSE-PhyloTree Docker and Singularity images, along with all scripts, test data, and documentation, are available at https://github.com/OcMalde/fuse-phylotree.

## Conclusion

FUSE-PhyloTree offers a pipeline for exploring sequence conservation modules and their association with functionnal annotations through phylogenomic analysis. This is the first tool to simultaneously exploit a new method of partial local multiple alignment, three-level phylogenetic reconciliation methods and ancestral reconstruction of protein functions. This comprehensive framework enables in-depth analysis of sequence-function relationships in multi-functional protein families. Ultimately, FUSE-PhyloTree predicts functional sequence regions, making it an invaluable asset for discovering new therapeutic targets. All these advanced methods are seamlessly integrated via custom Python scripts and can be deployed using containerized images for both Docker and Singularity.

## Acknowledgments

The authors thank the excellent support of the GenOuest bioinformatics core facility.

## Funding

This work was supported by Université de Rennes, INSERM and Inria.

## APPENDIX

### Illustrated use-case of FUSE-PhyloTree on the fibulin protein family

We present here a practical example of the application of FUSE-PhyloTree to a multi-domain, multifunctional protein family: the *fibulins* (3).

Fibulins is a family of extracellular matrix glycoproteins associated with basement membrane and elastic fibers. Eight fibulin paralogs are identified in humans and are characterized by variable lengths and domain compositions, including a fibulin-type C-terminal domain preceded by tandem calcium-binding epidermal growth factor (EGF)-like domains (9). Their expression is altered in many diseases such as cancer where they are thought to play both anti- and pro-oncogenic roles depending on their molecular interaction with other proteins (5). The identification of specific functional sequence regions within fibulins is therefore promising for the development of new targeted therapeutic strategies

In this appendix, we will consider the different proteinprotein interactions of the fibulins as functions of interest, and illustrate step-by-step how to use FUSE-PhyloTree in order to identify and explore the local sequence conservations that may be associated with these interactions.

#### Download

The first step is to install the tool. FUSE-PhyloTree is available as a Singularity or Docker image, which are available for download on our GitHub (https://github.com/OcMalde/fuse-phylotree).

We will use here Singularity (version 3.6.3). If not already available in your environment, Singularity will need to be installed (see https://docs.sylabs.io/guides/3.0/user-guide/installation.html).

The FUSE-PhyloTree image `fuse-phylotree.sif` can be downloaded by executing the following command in a terminal:

`singularity pull` https://github.com/OcMalde/fuse-phylotree/releases/download/V1.0.0/fuse-phylotree.sif

#### Input

Then we need to prepare the input data. The program takes three different files as input providing:

- A **set of reference sequences**, in Fasta format, containing the protein sequences of the family (orthologs and paralogs).
- To enable automatic retrieval of the species tree, FUSE-PhyloTree requires the Fasta identifiers to be in the form *>*`SeqID_taxid`, where `SeqID` is a unique identifier for the sequence, such as a RefSeq ID, and `taxid` is the taxonomic identifier of the organism in NCBI Taxonomy database. For studies targeting a family of human paralogs, FUSE-PhyloTree provides the helper script `make_orthogroup_fasta.sh` to automatically generate this formated Fasta file from a file containing the RefSeq ID of the sequences with pre-computed orthogroups for nine bilaterian species available in the FUSE-PhyloTree image. Using this script with the 8 human fibulins as input, after discarding a suspect short ortholog that does not contain the required fibulin domains, we obtained 59 fibulin sequences in 9 bilaterian species (see Table S1). These sequences, used hereafter as reference sequences, are available in the file `fibulin_59.fasta` (See GitHub for Files).
- A **rooted, binary phylogenetic tree**, in Newik format, of the set of reference sequences. For our study, we considered members of the LTBP family (latent transforming growth factor (TGF-*β*)-binding proteins) as close relatives of the fibulins (12) enabling to root the tree. The outgroup sequences consisted of LTBP-1 and LTBP-4 human paralogs and their orthologs from the eight other bilaterian species considered for the reference sequence file. Adding these sequences to the 59 fibulins, a phylogenetic tree was inferred. Its root was manually set and the outgroup was removed using iTOL (7). The resulting phylogenetic tree is available in the file `fibulin_tree_root.tree` (See GitHub for Files).
- A set of **functional annotations of reference sequences**, in comma-separated value text format (CSV), where each row contains the known functional annotation of interest of a reference sequence in the form: `SeqID,Annotation1|Annotation2|…|AnnotationN` with the first value consisting of the sequence identifier and the second value specifying the annotations of the sequence separated by the | symbol.

In our use-case, the considered functional annotation are the protein-protein interactions (PPIs) currently known for the human fibulins. They were retrieved through the Proteomics Standard Initiative Common Query Interface (PSICQUIC) webservice (1) and filtered to retain only 97 PPIs shared by at least two human fibulin paralogs (Fig. S1). The resulting functional annotation file is `ppi_shared.csv` (See GitHub for Files).

**Table S1.**
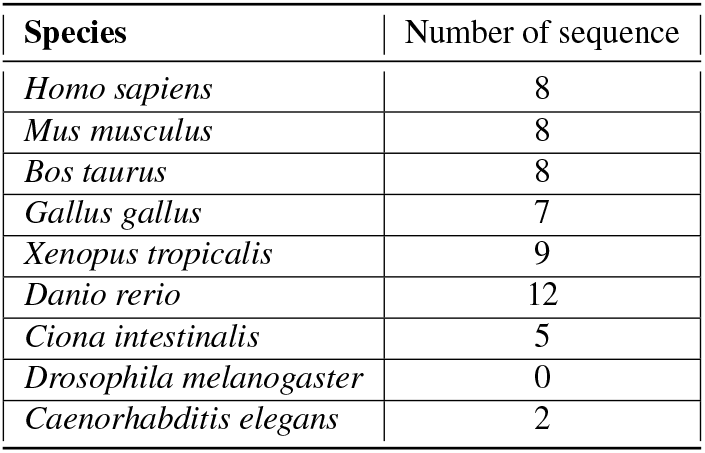
Number of fibulin paralogs per species.

| Species | Number of sequence |
| --- | --- |
| *Homo sapiens* | 8 |
| *Mus musculus* | 8 |
| *Bos taurus* | 8 |
| *Gallus gallus* | 7 |
| *Xenopus tropicalis* | 9 |
| *Danio rerio* | 12 |
| *Ciona intestinalis* | 5 |
| *Drosophila melanogaster* | 0 |
| *Caenorhabditis elegans* | 2 |

#### Run

The following bash script can be used to launch the analysis, using as inputs the three files presented above (“fibu-lin_59.fasta”, “ppi_shared.csv” and “fibulin_tree_root.tree”): #!/bin/bash

~~~
# File paths
image_path=“fuse-phylotree.sif”
file_fasta=“fibulin_59.fasta”
file_annotation=“ppi_shared.csv”
gene_tree=“fibulin_tree_root.tree” output_dir=“dir_analysis_fuse_phylotree”
# Command to launch tool
cmd=“python3 /fuse-phylotree/fuse-phylotree.py
--output_directory ${output_dir}
$file_fasta $file_annotation $gene_tree”
# Start analysis
singularity exec ${image_path} ${cmd}
~~~

where `dir_analysis_fuse_phylotree` specifies the directory where the intermediate files of the workflow are saved for advanced users.

#### Output

The main outputs are located in the working directory:

- The file named `0_gene.tree` (Newick format - See GitHub for Files) is the gene tree with the canonical renaming of genes used by FUSE-PhyloTree.
- The file named `1_modules_and_functions_evolution.csv` (See GitHub for Files) details the modules and functions assigned by FUSE-PhyloTree to each (ancestral or observed) gene, along with a view of the modules and functions lost or gained by the gene with respect to its closest ancestor in the tree.
- The file named `2_module_descriptions.csv` (See GitHub for Files) provides for each module the location (protein ID, start and end positions in the protein sequence) of its segments.
- Finally a directory named `3_visuReconc` (See GitHub for Files) gathers all the files required for viewing the genes, modules and functions evolution using iTOL.

In the end, in our 59 fibulins use case, the file `2_module_descriptions.csv` lists the 508 modules identified by FUSE-PhyloTree. Their inferred co-evolution with functions can be examined by looking to the file `1_modules_and_functions_evolution.csv`. It enables for instance to distinguish among the 116 ancestral genes of the phylogeny, 48 ancestral genes with a gain of modules and, among them, 13 ancestral genes with the coappearance of at least one module and one PPI (see Table S2). Among these 13 ancestral genes, 4 have at least two human descendants: G50, G110, G111 and G115. Note that, in addition to their importance in identifying relevant conserved modules, orthologs enable FUSE-PhyloTree to predict the co-emergence of modules and function of ancestral gene with a single human descendant, for instance the ancestral gene G94 of fibulin 4, which gains two PPIs, CREB5 and FBLN5 and 3 conserved modules B615, B648 and B651. Fig. S2 shows an iTOL visualization of the results, obtained from the files available in `3_visuReconc` directory. The interactive version of the tree is available at https://itol.embl.de/tree/13125423118208151734008923

#### Interpretation

To illustrate the exploration of the results, we propose to zoom in on the gene G111 from Table S2, the common ancestor of three human fibulins (FBLN3, FBLN4 and FBLN5) called the elastic fibulins.

G111 is predicted to gain 6 interactions (with DYRK1A, GFI1B, LTBP1, MEOX2, OTX1 and TGFB1), 3 of which (DYRK1A, LTBP1 and TGFB1) are specific to this sub-tree. Interestingly, LTBP1 and TGFB1 are biologically closely related, as LTPB1 is known to bind the latent form of TGFB1. It can be noted that DYRK1A has also connections to TGFB1, as it has been shown to interact with TGFB1 signaling pathway (2). Alongside its gained interactions, G111 is predicted to gain 10 modules (B77, B565, B595, B612, B622, B624,

B625, B626, B628 and B644) shown in Fig. S3. Let us remark that 8 of the gained modules are present in all the three sequences. The two other modules B77 and B644 are respectively absent in FBLN4 and FBLN3, but are respectively present in orthologs of FBLN4 (in *Xenopus tropicalis, Mus musculus* and *Bos taurus*) and FBLN3 (in *Gallus gallus* and *Mus musculus*), supporting FUSE-PhyloTree prediction that they were gained in G111 and subsequently lost in specific lineages.

The locations of the conserved segments of these modules in the sequences of G111’s human descendants are shown in Fig. S4 (see also Fig. S5).

**Fig. S1.**
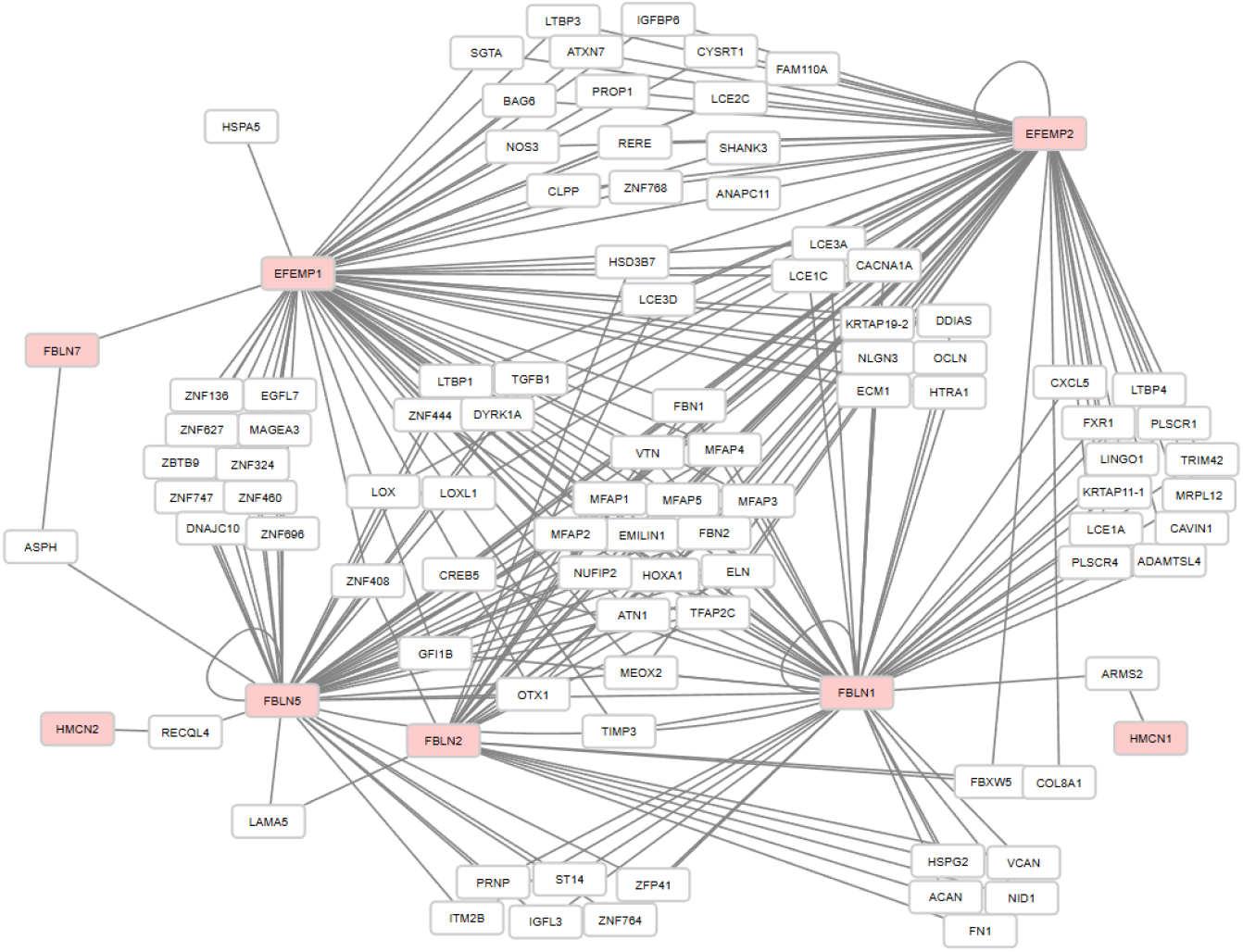
Network of protein-protein interactions (PPIs). Visualization was performed with the Cytoscape tool (13) and shows the 96 PPIs shared by at least two human fibulins (red nodes). *Legend* : HMCN1 : hemicentin-1, HMCN2 : hemicentin-2, FBLN2 : fibulin-2, FBLN1 : fibulin-1, FBLN7 : fibulin-7, FBLN5 : fibulin-5, FBLN4 : fibulin-4 (EFEMP2 gene) and FBLN3 : fibulin-3 (EFEMP1 gene). Hemicentins 1 and 2 are also known as fibulin 6 and 8 respectively.

The modules gained at G111 predominantly span the characteristic C-terminal domain of fibulins, detected by “Fibulin C-terminal Ig-like” PFAM model (PF22914, 10). Modules B624, B625 B626 and B628 cover this PFAM domain and enable to refine its functional role by revealing a specific conservation of the region in FBLN 3, 4 and 5 subtree that is probably linked to their specific interactions. This prediction was confirmed for LTBP1 by experimental studies showing the interaction of the C-terminal domain of FBLN4 and FBLN5 with LTBP proteins (6, 11). Additionally, module B624 extends beyond the identified PFAM domain, suggesting an expanded functional region of interest.

The remaining gained modules (B565, B595, B612, B644, B77 and B622) map to calcium-binding EGF-like (EGF_CA) domains of fibulins detected by SMART (SM0017, 8). Modules B565 and B595 are included in EFG_CA 1, an atypical EFG_CA domain (with two of the six conserved cysteine residues of the domain missing) that is nonetheless functionally important as shown by the study of the mutation of FBLN3 cysteine residue at position 55 (15), which is also found in B565. Modules B612, B644, B77, and B622 are found in EGF_CA domains 2, 4, 5, and 6, respectively. Interestingly, each is positioned around the last cysteine residue of the domain, in its exposed loop and second strand of C-terminal *β*-sheet region (see Fig S5). Combined with their residue composition shown in Fig. S3, in particular the presence of negatively charged conserved residues, this reinforces their potential role in interaction specificity and makes them interesting candidates for therapeutic targeting, which merits further investigation.

To conclude, analysis of the fibulin family using FUSE-PhyloTree revealed the evolutionary history of 508 conserved modules and 97 PPIs. It identified 13 ancestral genes for which conserved modules and PPI co-emerged. The interpretation of these predictions was illustrated for the ancestral gene G111, demonstrating that FUSE-PhyloTree could identify modules involved in experimentally validated interactions and also reveal novel potential functional modules. Beyond these direct predictions, FUSE-PhyloTree can also be used to interactively explore the evolution of functions and sequence conservation along the phylogenetic gene tree, and gain deeper insights into the family. For example, it would be interesting to explore with iTOL the direct ancestors and descendants of the gene G111 to refine predictions, or entire paths to G111, or from G111, to obtain information on the evolution of modules and their specificity.

All the files resulting from the analysis are available on GitHub at (https://github.com/OcMalde/fuse-phylotree/tree/main/data/analyse_fibulin/run_singularity_fibulin).

**Fig. S2.**
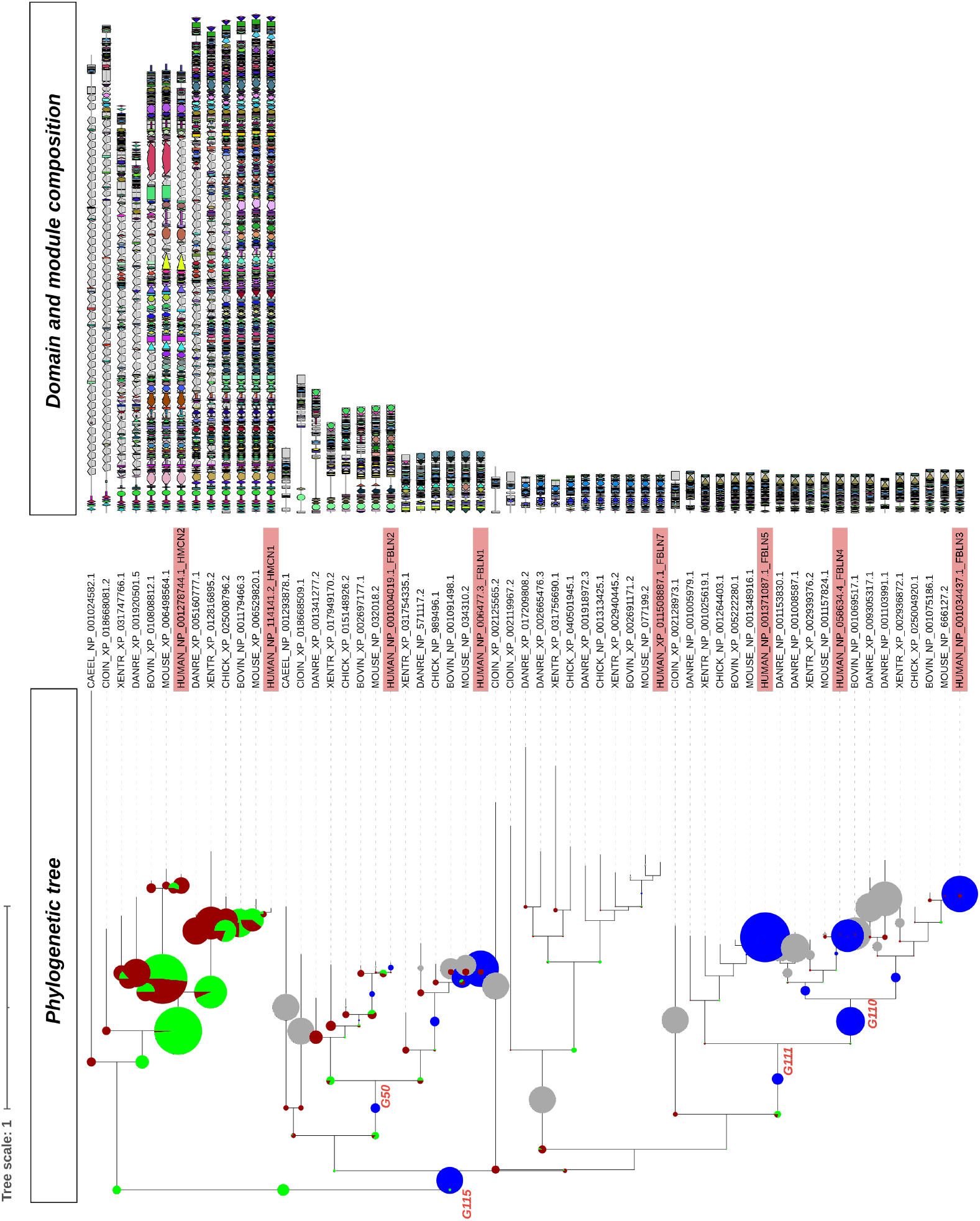
Representation on a phylogenetic tree of the modules and PPIs emergence. The phylogenetic tree shows the number of gains and losses of PPIs (blue and grey circles, respectively) and modules (green and red respectively in the pie chart). The leaf labels of the human fibulins are highlighted in red and followed by the abbreviation of the paralog name. In the domain and module composition section, each module is represented by a unique combination of shape and color. The grey shapes in the background indicate the location of Pfam domains. *Legend* : CAEEL : *Caenorhabditis elegans*, CIOIN : *Ciona intestinalis*, DANRE : *Danio rerio*, XENTR : *Xenopus tropicalis*, CHICK : *Gallus gallus*, BOVIN : *Bos taurus*, MOUSE : *Mus musculus*, HUMAN : *Homo sapiens*, HMCN1 : hemicentin-1, HMCN2 : hemicentin-2, FBLN2 : fibulin-2, FBLN1 : fibulin-1, FBLN7 : fibulin-7, FBLN5 : fibulin-5, FBLN4 : fibulin-4 (EFEMP2 gene) et FBLN3 : fibulin-3 (EFEMP1 gene). Hemicentins 1 and 2 are also known as fibulin 6 and 8 respectively. The presence of large red and green pie charts in the upper part of the tree can be explained by the length of the proteins involved, the hemicentins, compared with the other proteins in the tree. Their length implies a significant gain in modules. The presence of blue and grey pie charts towards the bottom (fibulin 3, 4 and 5) and in the center (fibulin 1) is understandable given the large number of PPIs known for these human proteins.

**Table S2.**
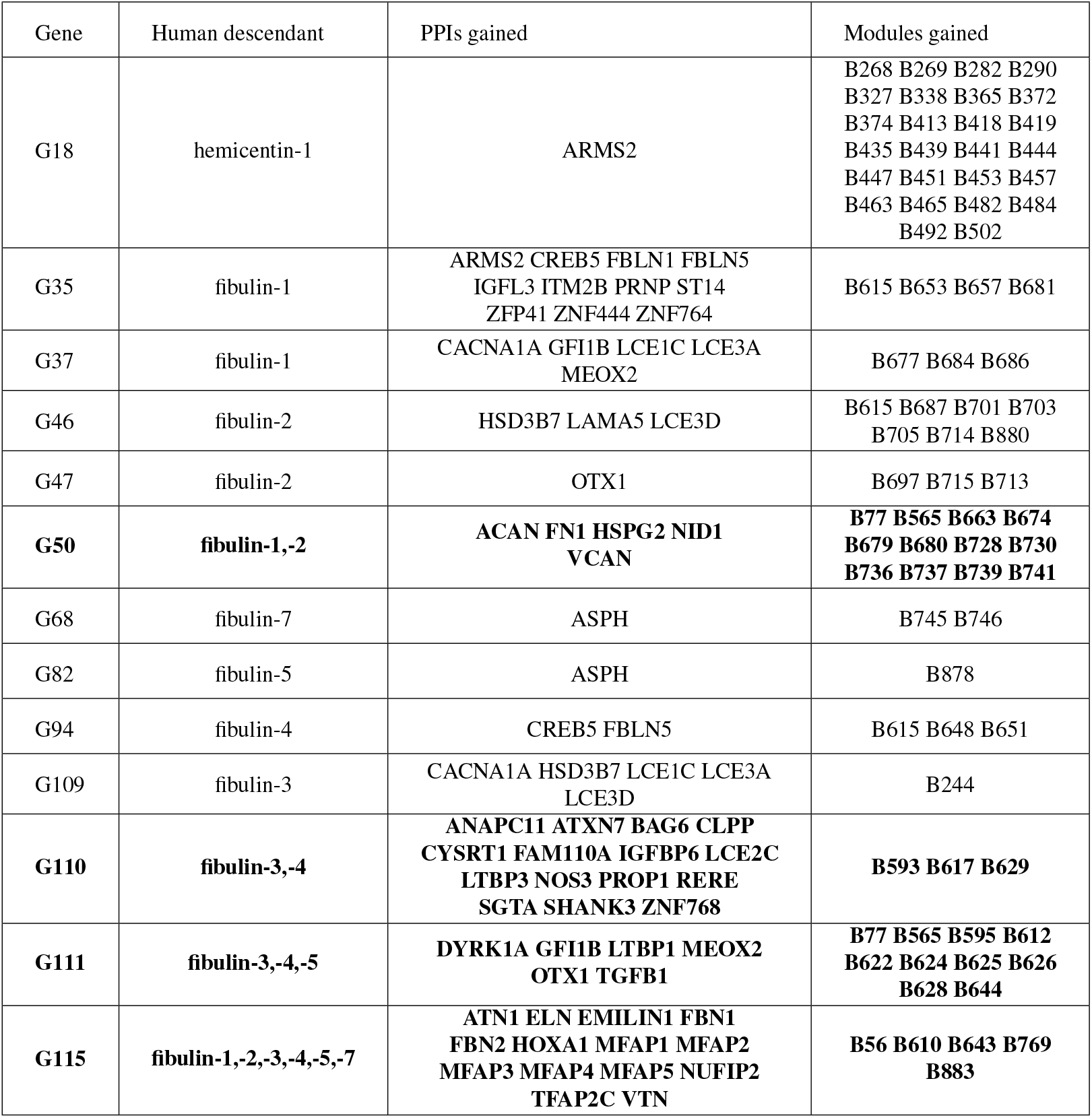
Table of 13 nodes with module-PPI co-emergence. For each node, its (i) name (Gene), (ii) human descendants, (iii) gained PPI and (iv) gained modules are given. Ancestral nodes highlighted in bold have at least two human paralogs as descendants.

**Fig. S3.**
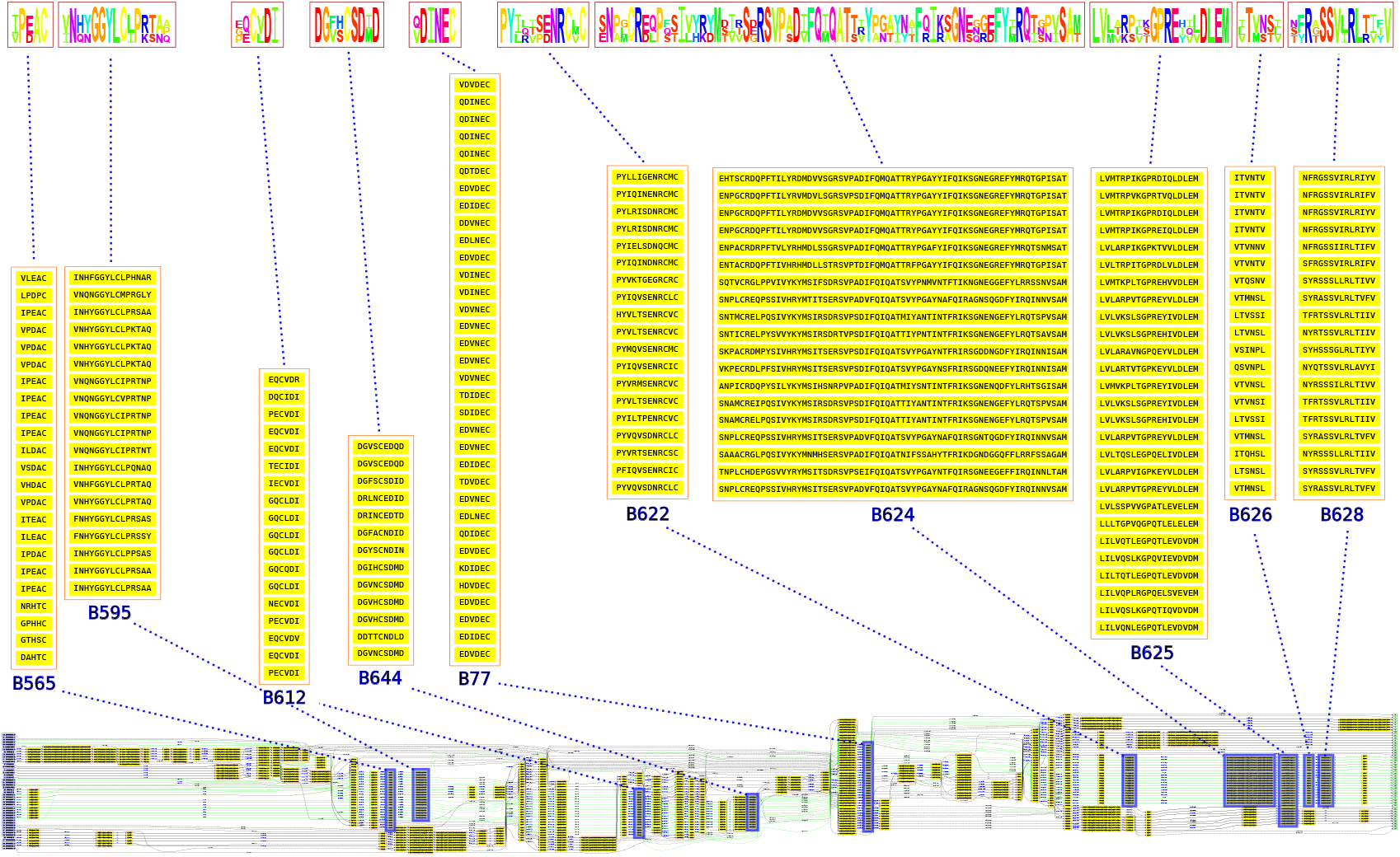
Modules gained at G111 on fibulins. Hemicentin sequences being too long to display, only fibulin sequences from G115 subtree are considered here. The partial local multiple alignment (PLMA) of the sequences used to identify the conserved modules is shown at the bottom of the figure. Set of conserved segments are highlighted in yellow and surrounded by red boxes. Many conserved segment sets involve only a few sequences, but only those involving at least 5 segments are displayed here. Lines indicate the path of each sequence between its conserved segments, green color being used for descendant sequences from G111. Modules predicted to be gained at ancestor gene G111 (B565, B595, B612, B644, B77, B622, B624, B625, B626 and B628) are highlighted in blue in the PLMA with a zoom on their segment content above and, on the top of the figure, a sequence logo (4) showing their positional residue conservation.

**Fig. S4.**
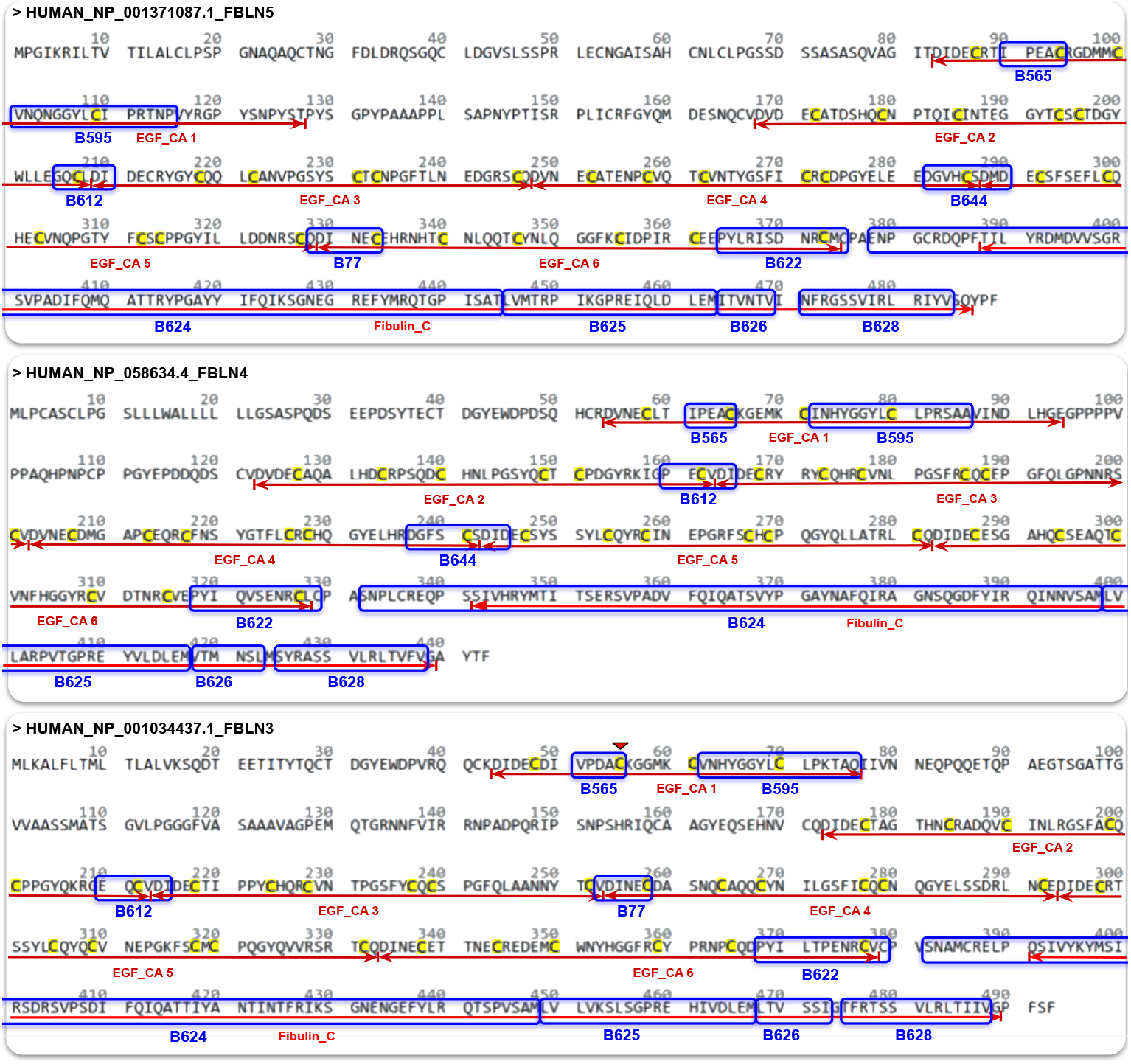
Position of modules gained by the G111 ancestral gene in its human descendants. Segments of modules gained at G111 (B565, B595, B612, B644, B77, B622, B624, B625, B626 and B628) are shown here circled by blue boxes, with their name underneath. Note that modules B77 and B644 are not conserved in these FBLN4 and FBLN3 sequences respectively. The six occurrences of SMART SM0017 domain (EGF_CA), numbered from 1 to 6 in each sequence) and the occurrence of PFAM PF22914 domain (Fibulin_C) in the sequences are underlined by red arrows, with their name underneath. Conserved cysteine residues in EGF_CA domains are highlighted in yellow and FBLN3’s cysteine residue at position 55 is marked by red triangle.

**Fig. S5.**
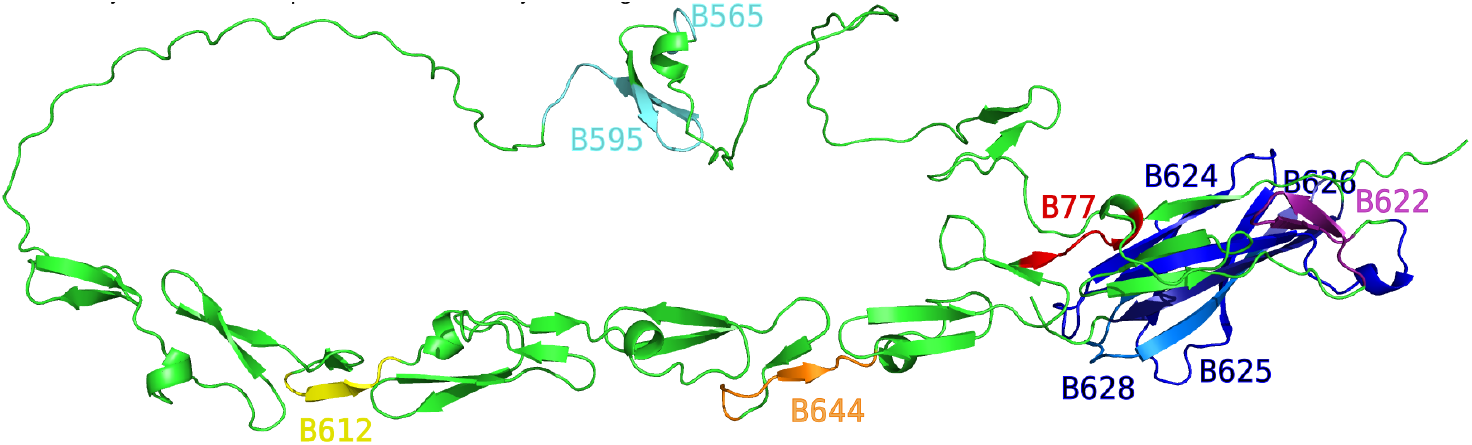
Location of G111 gained modules on predicted 3D structure of HUMAN_NP_001371087.1_FBLN5 by AlphaFold (AF-G3XA98-F1-v4, 14).

## Notes

https://github.com/OcMalde/fuse-phylotree

### Competing Interest Statement

The authors have declared no competing interest.

## References

1 Salvador Capella-Gutiérrez, José M. Silla-Martínez, and Toni Gabaldón. trimal: a tool for automated alignment trimming in large-scale phylogenetic analyses. Bioinformatics, 25(15): 1972–1973, June 2009. ISSN 1367-4803.

2 Olivier Dennler, François Coste, Samuel Blanquart, Catherine Belleannée, and Nathalie Théret. Phylogenetic inference of the emergence of sequence modules and protein-protein interactions in the adamts-tsl family. PLOS Computational Biology, 19(8):e1011404, August 2023. ISSN 1553-7358.

3 Robert C. Edgar. Muscle5: High-accuracy alignment ensembles enable unbiased assessments of sequence homology and phylogeny. Nature Communications, 13(1), November 2022. ISSN 2041-1723.

4 Jonathan A. Eisen. Phylogenomics: Improving functional predictions for uncharacterized genes by evolutionary analysis. Genome Research, 8(3):163–167, March 1998. ISSN 1549-5469.

5 Stéphane Guindon, Jean-François Dufayard, Vincent Lefort, Maria Anisimova, Wim Hordijk, and Olivier Gascuel. New algorithms and methods to estimate maximum-likelihood phylogenies: Assessing the performance of phyml 3.0. Systematic Biology, 59(3):307–321, March 2010. ISSN 1063-5157.

6 Sohta A Ishikawa, Anna Zhukova, Wataru Iwasaki, and Olivier Gascuel. A fast likelihood method to reconstruct and visualize ancestral scenarios. Molecular Biology and Evolution, 36(9):2069–2085, May 2019. ISSN 1537-1719.

7 Goulven Kerbellec. Apprentissage d’automates modélisant des familles de séquences pro-téiques (Learning automata modelling families of protein sequences). Theses, Université Rennes 1, June 2008.

8 Ivica Letunic and Peer Bork. Interactive tree of life (itol) v6: recent updates to the phylo-genetic tree display and annotation tool. Nucleic Acids Research, 52(W1):W78–W82, April 2024. ISSN 1362-4962.

9 Lei Li and Mukul S. Bansal. Simultaneous Multi-Domain-Multi-Gene Reconciliation Under the Domain-Gene-Species Reconciliation Model, page 73–86. Springer International Publishing, 2019. ISBN 9783030202422.

10 Typhaine Paysan-Lafosse, Matthias Blum, Sara Chuguransky, Tiago Grego, Beatriz Lázaro Pinto, Gustavo A Salazar, Maxwell L Bileschi, Peer Bork, Alan Bridge, Lucy Colwell, Julian Gough, Daniel H Haft, Ivica Letunić, Aron Marchler-Bauer, Huaiyu Mi, Darren A Natale, Christine A Orengo, Arun P Pandurangan, Catherine Rivoire, Christian J A Sigrist, Ian Sillitoe, Narmada Thanki, Paul D Thomas, Silvio C E Tosatto, Cathy H Wu, and Alex Bateman. Interpro in 2022. Nucleic Acids Research, 51(D1):D418–D427, November 2022. ISSN 1362-4962.

11 Clemens Rauer, Neeladri Sen, Vaishali P. Waman, Mahnaz Abbasian, and Christine A. Orengo. Computational approaches to predict protein functional families and functional sites. Current Opinion in Structural Biology, 70:108–122, October 2021. ISSN 0959-440X.

12 Yan Wang, Hang Zhang, Haolin Zhong, and Zhidong Xue. Protein domain identification methods and online resources. Computational and Structural Biotechnology Journal, 19: 1145–1153, 2021. ISSN 2001-0370.

13 Yi-Chieh Wu, Matthew D. Rasmussen, Mukul S. Bansal, and Manolis Kellis. Treefix: Statistically informed gene tree error correction using species trees. Systematic Biology, 62(1): 110–120, November 2012. ISSN 1063-5157.

## References for Appendix

1. Bruno Aranda, Hagen Blankenburg, Samuel Kerrien, Fiona S. L. Brinkman, Arnaud Ceol, Emilie Chautard, Jose M. Dana, Javier De Las Rivas, Marine Dumousseau, Eugenia Galeota, Anna Gaulton, Johannes Goll, Robert E. W. Hancock, Ruth Isserlin, Rafael C. Jimenez, Jules Kerssemakers, Jyoti Khadake, David J. Lynn, Magali Michaut, Gavin O’Kelly, Keiichiro Ono, Sandra Orchard, Carlos Prieto, Sabry Razick, Olga Rigina, Lukasz Salwinski, Milan Simonovic, Sameer Velankar, Andrew Winter, Guanming Wu, Gary D. Bader, Gianni Cesareni, Ian M. Donaldson, David Eisenberg, Gerard J. Kleywegt, John Overington, Sylvie Ricard-Blum, Mike Tyers, Mario Albrecht, and Henning Hermjakob. PSICQUIC and PSIS-CORE: Accessing and scoring molecular interactions. Nature Methods, 8(7):528–529, July 2011. ISSN 1548-7105.

2. Ying Cao, Ruolan Qian, Ruilian Yao, Quan Zheng, Chen Yang, Xupeng Yang, Shuyi Ji, Linmen Zhang, Shujie Zhan, Yiping Wang, Tianshi Wang, Hui Wang, Chun-Ming Wong, Shengxian Yuan, Christopher Heeschen, Qiang Gao, René Bernards Wenxin Qin, and Cun Wang. DYRK1A-TGF-β signaling axis determines sensitivity to OXPHOS inhibition in hepatocellular carcinoma. Developmental Cell, pages S1534–5807(24)00775–5, January 2025. ISSN 1878-1551.

3. Marion A. Cooley and W. Scott Argraves. The Fibulins. In Robert P. Mecham, editor, The Extracellular Matrix: An Overview, pages 337–367. Springer, Berlin, Heidelberg, 2011. ISBN 978-3-642-16555-9.

4. Gavin E. Crooks, Gary Hon, John-Marc Chandonia, and Steven E. Brenner. Weblogo: A sequence logo generator. Genome Research, 14(6):1188–1190, 2004.

5. William M. Gallagher, Caroline A. Currid, and Linda C. Whelan. Fibulins and cancer: Friend or foe? Trends in Molecular Medicine, 11(7):336–340, July 2005. ISSN 1471-4914.

6. Maretoshi Hirai, Masahito Horiguchi, Tetsuya Ohbayashi, Toru Kita, Kenneth R. Chien, and Tomoyuki Nakamura. Latent TGF-beta-binding protein 2 binds to DANCE/fibulin-5 and regulates elastic fiber assembly. The EMBO journal, 26(14):3283–3295, July 2007. ISSN 0261-4189.

7. Ivica Letunic and Peer Bork. Interactive Tree of Life (iTOL) v6: Recent updates to the phylogenetic tree display and annotation tool. Nucleic Acids Research, 52(W1):W78–W82, July 2024. ISSN 0305-1048.

8. Ivica Letunic, Supriya Khedkar, and Peer Bork. Smart: recent updates, new developments and status in 2020. Nucleic Acids Research, 49(D1):D458–D460, 10 2020. ISSN 0305-1048.

9. Deviyani Mahajan, Sudhakar Kancharla, Prachetha Kolli, Amarish Kumar Sharma, Sanjeev Singh, Sudarshan Kumar, Ashok Kumar Mohanty, and Manoj Kumar Jena. Role of Fibulins in Embryonic Stage Development and Their Involvement in Various Diseases. Biomolecules, 11(5):685, May 2021. ISSN 2218-273X.

10. Jaina Mistry, Sara Chuguransky, Lowri Williams, Matloob Qureshi, Gustavo A Salazar, Erik L L Sonnhammer, Silvio C E Tosatto, Lisanna Paladin, Shriya Raj, Lorna J Richardson, Robert D Finn, and Alex Bateman. Pfam: The protein families database in 2021. Nucleic Acids Research, 49(D1):D412–D419, 10 2020. ISSN 0305-1048.

11. Kazuo Noda, Branka Dabovic, Kyoko Takagi, Tadashi Inoue, Masahito Horiguchi, Maretoshi Hirai, Yusuke Fujikawa, Tomoya O. Akama, Kenji Kusumoto, Lior Zilberberg, Lynn Y. Sakai, Katri Koli, Motoko Naitoh, Harald von Melchner, Shigehiko Suzuki, Daniel B. Rifkin, and Tomoyuki Nakamura. Latent TGF-β binding protein 4 promotes elastic fiber assembly by interacting with fibulin-5. Proceedings of the National Academy of Sciences of the United States of America, 110(8):2852–2857, February 2013. ISSN 1091-6490.

12. Ian B. Robertson, Masahito Horiguchi, Lior Zilberberg, Branka Dabovic, Krassimira Hadjiolova, and Daniel B. Rifkin. Latent TGF-β-binding proteins. Matrix Biology: Journal of the International Society for Matrix Biology, 47:44–53, September 2015. ISSN 1569-1802.

13. Paul Shannon, Andrew Markiel, Owen Ozier, Nitin S. Baliga, Jonathan T. Wang, Daniel Ramage, Nada Amin, Benno Schwikowski, and Trey Ideker. Cytoscape: A Software Environment for Integrated Models of Biomolecular Interaction Networks. Genome Research, 13(11):2498–2504, November 2003. ISSN 1088-9051, 1549-5469.

14. Mihaly Varadi, Damian Bertoni, Paulyna Magana, Urmila Paramval, Ivanna Pidruchna, Malarvizhi Radhakrishnan, Maxim Tsenkov, Sreenath Nair, Milot Mirdita, Jingi Yeo, Oleg Kovalevskiy, Kathryn Tunyasuvunakool, Agata Laydon, Augustin Žídek, Hamish Tomlinson, Dhavanthi Hariharan, Josh Abrahamson, Tim Green, John Jumper, Ewan Birney, Martin Steinegger, Demis Hassabis, and Sameer Velankar. Alphafold protein structure database in 2024: providing structure coverage for over 214 million protein sequences. Nucleic Acids Research, 52(D1):D368–D375, 11 2023. ISSN 0305-1048.

15. DaNae R. Woodard, Steffi Daniel, Emi Nakahara, Ali Abbas, Sophia M. DiCesare, Gracen E. Collier, and John D. Hulleman. A loss-of-function cysteine mutant in fibulin-3 (efemp1) forms aberrant extracellular disulfide-linked homodimers and alters extracellular matrix composition. Human Mutation, 43(12):1945–1955, 2022.

